# Efficient test for deviation from Hardy Weinberg Equilibrium with known or ambiguous typing in highly polymorphic loci

**DOI:** 10.1101/2024.03.19.585658

**Authors:** Or Shkuri, Sapir Israeli, Yuli Tshuva, Martin Maiers, Yoram Louzoun

## Abstract

The Hardy-Weinberg Equilibrium (HWE) assumption is essential to many population genetics models. Multiple tests were developed to test its applicability in observed genotypes. Current methods are divided into exact tests applicable to small populations and a small number of alleles, and approximate goodness of fit tests. Existing tests cannot handle ambiguous typing in multi-allelic loci. We here present a novel exact test (UMAT - Unambiguous Multi Allelic Test) practically not limited in the number of alleles and population size, based on a perturbative approach around the current observations. We show its accuracy in the detection of deviation from HWE. We then propose an additional model to handle ambiguous typing using either sampling into UMAT or a goodness of fit test with a variance estimate taking ambiguity into account, named ASTA (Asymptotic Statistical Test with Ambiguity). We show the accuracy of ASTA and the possibility to detect of the source of deviation from HWE. We apply these tests to the HLA loci to recover multiple previously reported deviations from HWE, and a large number of new ones.

## Introduction

One of the most frequent assumptions regarding the allele pair compositions in a well-mixed populations population is the Hardy-Weinberg equilibrium (HWE). The HWE presumes random-pairing of alleles in the population. Formally, one assumes constant genetic variation in a given locus, and that the allele probability in the two chromosomes of each host are pairwise independent. The HWE has been tested in many settings [14], including among many others SNPs [28], multi-allele loci, homogeneous vs structured populations [13], polyploid [31] vs diploids. We here focus on a single polymorphic locus. A classical example of such a locus would be the human chromosome 6 HLA region[1]. The HWE assumption in HLA-based population genetics has been tested in hundreds of studies [29]. In this case, given a *N* size sample from a population of size *N*_*P*_ with alleles taken from {1, …, *N*_*A*_} and unknown probabilities for the allele *i* - *p*(*i*), and for the pair {*i, j*} - *p*(*i, j*), the HWE assumption is that:

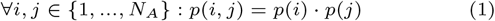

Note that one typically only computes the symmetric probabilities. In such a case, one would mark:

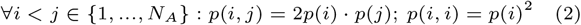

While *p*(*i, j*) is usually unknown, in most cases, one assumes that the observed allele pairs frequencies *O*(*i, j*) are a multinomial variable with the appropriate *p*(*i, j*). This is often further approximated when *N* is large by a binomial distribution for each *O*(*i, j*) using the appropriate *p*(*i, j*). However, in some cases, one does not know for a given individual *k* its precise allele pair. Instead, one may be given for the individual *k* a distribution of possible candidate pairs, each with an estimated probability in this sample of *O*_*k*_(*i, j*). One can then define the typing resolution score (TRS) [25]:

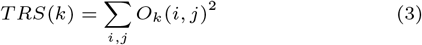

In the absence of ambiguity, *TRS*(*k*) = 1, else *TRS*(*k*) *<* 1. Note that in this context, the distribution of:

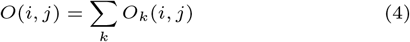

is not multinomial anymore, and for a high enough ambiguity, and large enough *N*, the central limit theorem holds for each *i, j*, and *O*(*i, j*) is normally distributed, even if it is small.

Multiple methods were developed to test the deviation from HWE. Those can be divided into a few categories. The first is the large-sample size goodness-of-fit tests. These tests assume a large population and therefore assume normality of *O*(*i, j*).

A very fundamental test in this group is the Chi-Square test [27]. Many variations of the Chi-Square were developed [6]. Such tests are only applicable to large populations.

An alternative is exact tests, based on a multinomial distribution assumption [21; 3; 12]. Such tests are more accurate than goodness of fit tests for small populations but require extensive computing [11]. A newer exact test was introduced [33] which works more efficiently and is therefore applicable for larger populations, but it is only applicable to 2 alleles.

A third group is likelihood tests. One example can be found in Elston et al [5] that works only for small samples. Another such test was proposed by Yu et al [34] that assumes normality and therefore is not as accurate as the exact tests for small samples. Other alternatives are Bayesian tests [22; 24] and Monte-Carlo-based methods [11; 20].

Recently, Montoya-Delgado et al. [8] stated that current exact tests are not feasible for more than three alleles. They introduced a recursion-based exact test that avoids repeating calculations. However, even this test is computationally expensive for more than four alleles. To deal with more than four alleles, they used a permutation test for an approximate p-value. We follow a part of their logic, since we also include permutations, and propose, using such permutations in a test that is not limited by the number of alleles.

None of the tests above can account for ambiguity in the observations in multi-allelic loci, as will be further defined. Recent papers aim to solve the problem of ambiguity [30; 19]. However, all the proposed tests with ambiguity are limited to bi-allelic systems.

To summarize, Goodness of fit tests require large populations, and exact tests are limited to small populations and a limited number of alleles. Beyond that, all existing methods cannot handle ambiguity in highly polymorphic loci. We propose two tests to solve for a single multi-allelic locus in diploids the two problems above. A) The high cost of exact tests for un-ambiguous samples. B) the variance estimate in goodness of fit tests in ambiguous samples.

To understand the effect of ambiguous typing, let us assume a set of 2 alleles: *a* and *A*, with equal probability. The homozygote probability *P* (*AA*) = 0.25, and in a sample of *N* alleles, the expected value of *O*(*AA*) is 0.25*N* and the variance is 0.25 *·* 0.75 *· N*. Now, assume instead that for each individual, we do not know the typing and assign *O*_*k*_(*AA*) = 0.25. In such a case, the expected value of *O*(*AA*) is still 0.25*N*. However, the variance is 0 (since all samples have *O*_*k*_(*AA*) = 0.25). One could propose to replace the unknown value, with a Bernoulli experiment, with the same probability, or produce a better estimate of the variance for the Wald test. We will show that this leads to different results.

## Materials and methods

### Allele pair simulation

Let us first assume that we observe the real allele pairs, (i.e., {*O*(*i, j*)} are known). We choose a value 0 ≤ *α* ≤ 1 that represents the fit of the simulation to the HWE.

- Produce a distribution {*p*(*i*)}_*i*_ s.t. 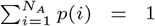 by generating a sequence of *q*_*i*_ ∼ *Uniform*(0, 2) for each allele and then normalizing the probability using a Softmax [2]: *P* = *Softmax*(*Q*). To compute the allele pairs probabilities, we first compute the probabilities assuming HWE:
- Calculate: *∀i, j* ∈ {1, 2, …, *N*_*A*_}:

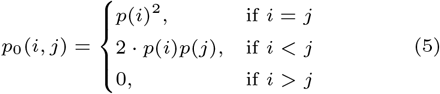
- In the case of deviation from HWE, we add the deviation and renormalize: *∀i, j* ∈ {1, 2,, …, *N*_*A*_}:

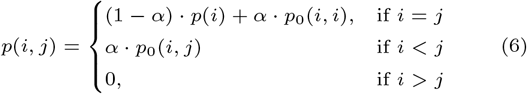

Finally,

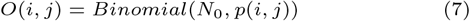

For the ambiguous case, we do not compute the binomial random variables. Instead, we compute a set of allele pairs with different probabilities for each sample. For each sample *k* with real alleles {*i, j*}, with probability *ρ*, we replace that by a group of all possible allele pairs, each with a probability {*p*(*t, m*)}_*t≤m*_, as defined in Eq (6).

### HLA genotype data

We studied 8,078,224 donors from the National Marrow Donor Program (NMDP) registry [10]. The dataset provided for each donor an HLA typing and the 21 detailed populations that combine into 5 broad populations (S1 Table) if known, any other typed race was converted to unknown. Each donor was imputed to produce the 20 most likely 5-locus haplotype pair using MR-GRIMM [15]. We computed for each donor its TRS.

### SNP data

We analyzed a dataset consisting of 13,258 Single Nucleotide Polymorphisms (SNPs) located on chromosome 6 from the thousand genome project [7]. Each of these SNPs can be represented by one of only two possible nucleotides. Our dataset comprises genetic information from 986 individuals.

### UMAT (Unambiguous Multi Allelic Test) Test for certain alleles

The UMAT test estimates the change in the log-likelihood following the random swap of 2 alleles toward HWE. These swaps are performed so that the marginal distribution is practically not changed along the analysis (to be precise, there are at most 4 alleles for which there is a deviation of at most a single individual from the marginal distributions). We then test how often the swapped distribution has a lower probability than the current distribution, only after a certain amount of swaps since the first swaps are biased. In the results presented here, we set the start to be 30, 000 swaps and use a total of 100, 000 iterations (in each iteration 2 swaps are performed), however, these values can be chosen arbitrarily.

#### Algorithm 1

Umat

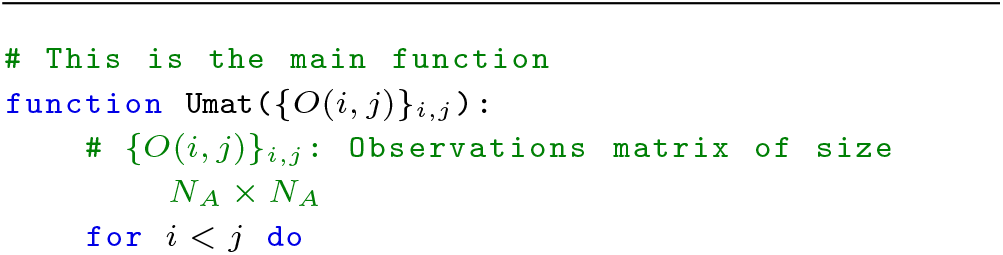
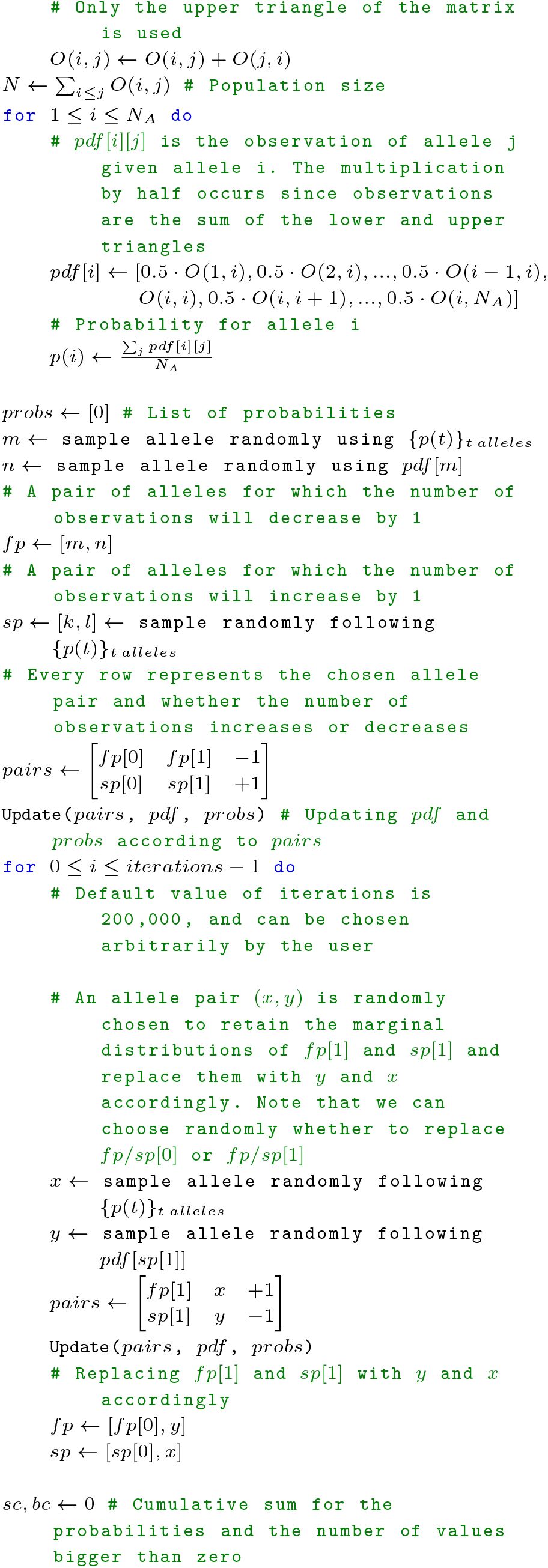
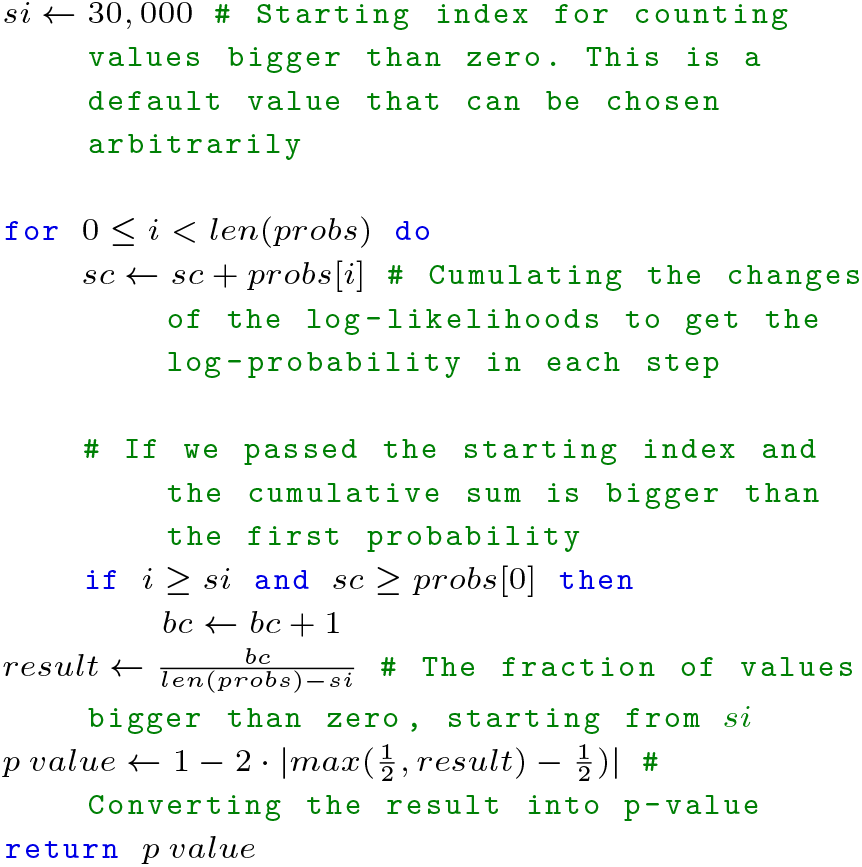

#### Algorithm 2

Function Update - UMAT

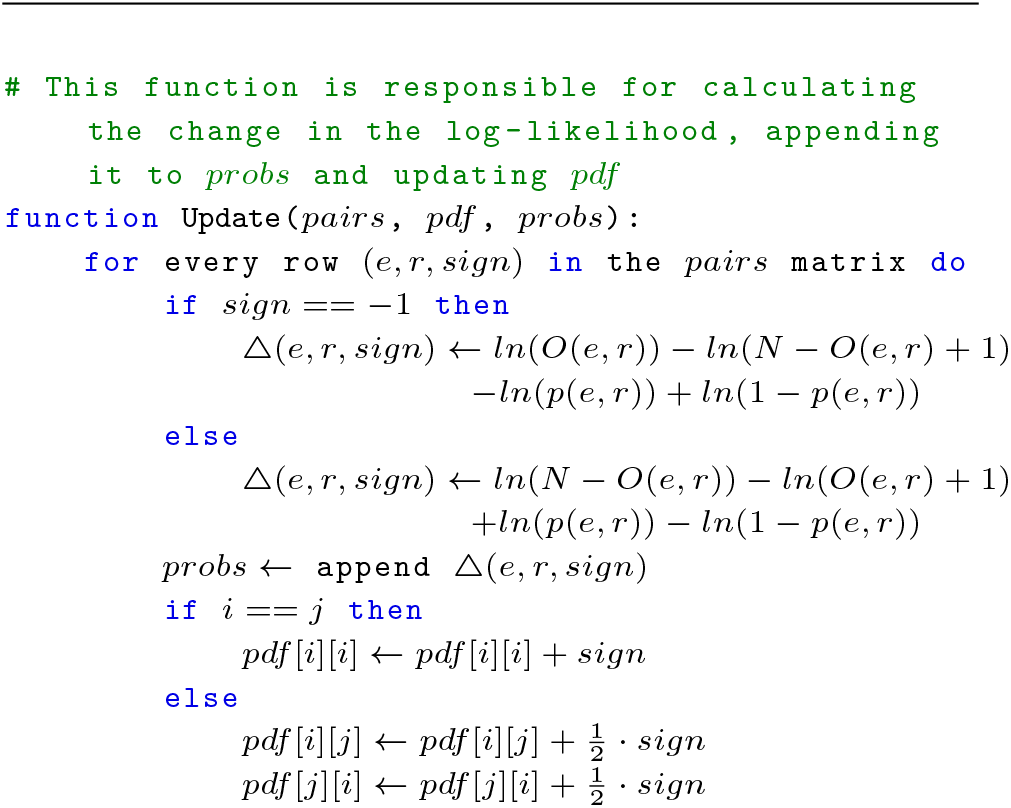

### ASTA (Asymptotic Statistical Test with Ambiguity)

Assuming ambiguous typing, one can adopt two options. A) Perform a sample following the probabilities of each pair in each donor and use the UMAT test above. B) Include in the goodness of fit test the ambiguity. The resulting algorithm is detailed below:

#### Algorithm 3

ASTA

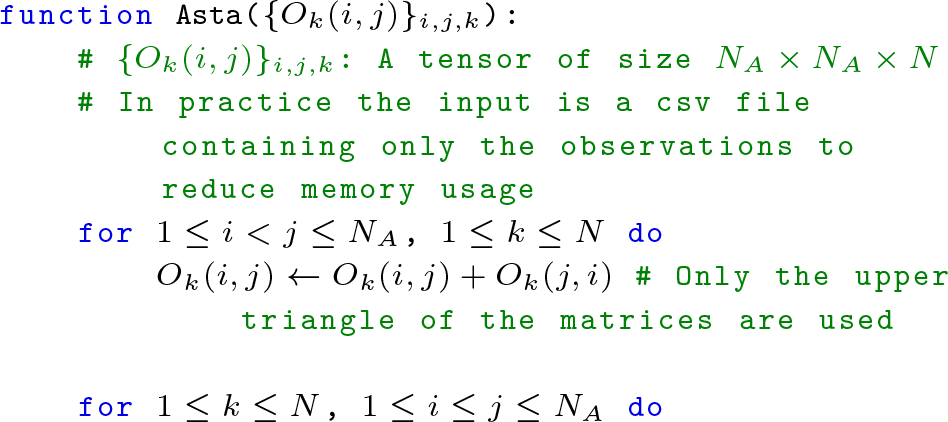
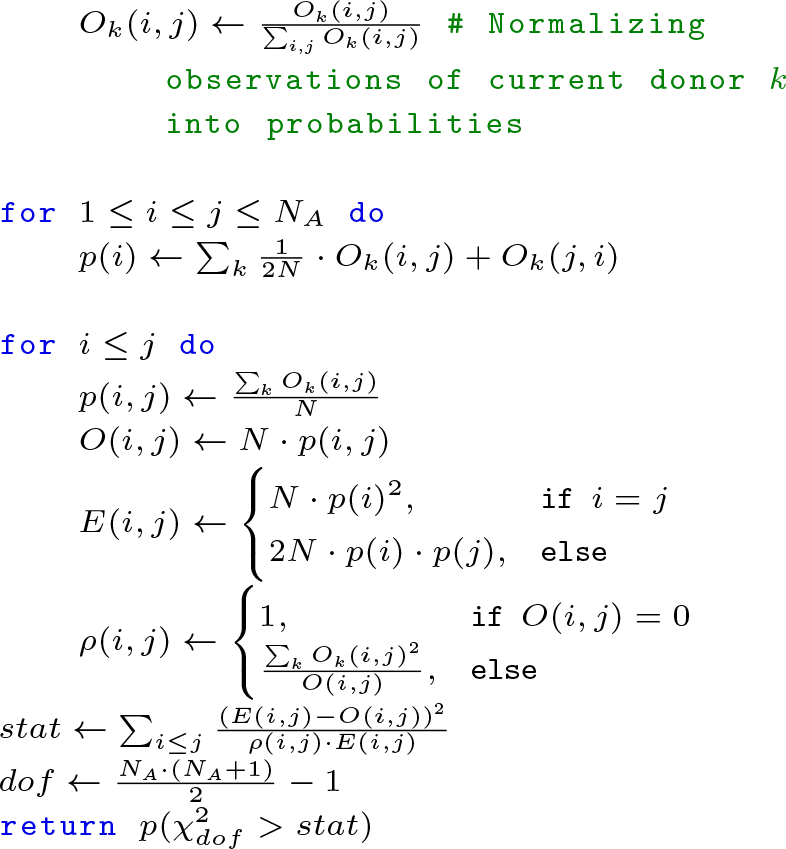

### Code and server

The methods that were introduced in this paper can be accessed via our python package https://github.com/louzounlab/HWE_Tests, including a user-friendly website that runs this package hwetests.math.biu.ac.il. Moreover, the code for all the simulations can be found in https://github.com/louzounlab/HWE_Simulations.

## Results

### Estimate of expected distribution

We follow a single highly polymorphic locus with a large number of candidate alleles, or similarly multiple phased loci with a large number of haplotypes. For the sake of notation, we denoted both haplotype and alleles as alleles, since do not test for Linkage Disequilibrium (LD), but only for HWE. The observations *O*_*k*_(*i, j*) are the probability that sample *k* has alleles *i* and *j*, where we further define *O*(*i, j*) = ∑_*k*_ *O*_*k*_(*i, j*).

In the absence of ambiguity, each sample has a single pair of alleles. In such a case, if the HWE assumption holds, *O*(*i, j*) can be approximated by a binomial Random Variable (RV) with an expected value and variance of *N · p*(*i, j*), where *N* is the sample size, for small enough *p*(*i, j*) (otherwise the variance is slightly smaller). If on the other hand, the ambiguity is large, and for each pair (*i, j*), *O*(*i, j*) is the sum of a large number of low probabilities, the central limit theorem holds, and *O*_*i,j*_ has approximately a normal distribution. We denoted the first case, the deterministic case (DC), and the second case the ambiguous case (AC).

In both cases, one needs an estimate of the expected *p*(*i, j*), assuming HWE. However, we only observe {*O*_*k*_(*i, j*)}_*i,j,k*_. We assumed the sample to be large enough to estimate the marginal distributions, with negligible errors:

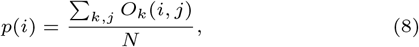

where *N* is the total number of samples. This assumption holds even for small *N*, as long as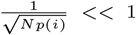. We then followed Eq 2 to compute the expected *p*(*i, j*), where here and all along, *p*(*i, j*) represents the expected pair distribution given the observed marginal distributions *p*(*i*) and HWE. In both AC and DC cases, the expected value of *O*(*i, j*) is *N · p*(*i, j*). However, as we will further show, the expected variance in AC is much lower than in DC.

### Gibbs sampling based exact test for DC case - UMAT

In the DC case, one can compute the observed sample probability as:

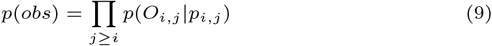

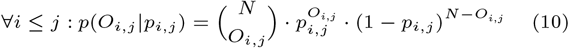

At this stage, instead of computing the entire distribution, we proposed to test the effect on the log-likelihood of performing a small perturbation, updating *O*(*k, l*) to *O*(*k, l*)+1, assuming the change is small enough that its effect on the marginal distributions *p*(*i*) (and thus on the expected *p*(*i, j*)) is negligible. The log ratio of the old to the new probabilities is:

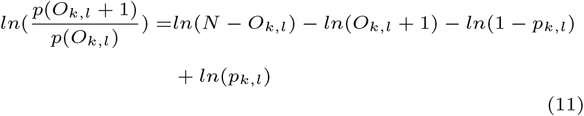

Similarly, for the reduction of 1 from *O*(*k, l*):

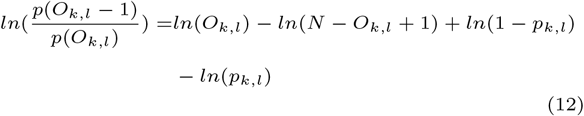

Following an addition and a removal operation, the total sample size does not change. However, the marginal distributions may change. We thus proposed a sampling strategy that would ensure the marginal distribution is not changed along iterations of additional and removal. At each iteration, except for the first and last one, UMAT adds and removes a single individual, maintaining the marginal distribution, up to two samples along the entire analysis, and ensuring the HWE in the change (Fig 1 A). In short (see methods), we keep a pair of alleles where an individual was removed, with two initial values *m, n* and a pair of alleles where an individual was added *k, l*. We choose an allele from the second pair (say *l*), and an additional allele chosen according to the distribution of *O*(*t, l*) *∀t* (i.e. all the observed allele pairs with allele *l*) and remove an individual and add it to the allele pair *m* (or *n*) and an allele chosen according to *p*(*m*). The new individual was added following the HWE assumption, and there are still only two alleles with one missing individual and two alleles with one additional individual. We then compute the increase/decrease in the log-likelihood (Eq 11, 12) to produce the total change in the log-likelihood from the initial stage following each change. Note that at no stage, do we compute the full probability distribution, only the changes. Finally, we check the fraction of changes where the cumulative change is larger than 0. If the HWE holds, this is a random walk around 0, and one can expect a distribution of the fraction of likelihoods above or below the initial values around 0.5. Otherwise, changes toward HWE should increase the likelihood. As such, most of the values will be above 0.

**Fig. 1.**
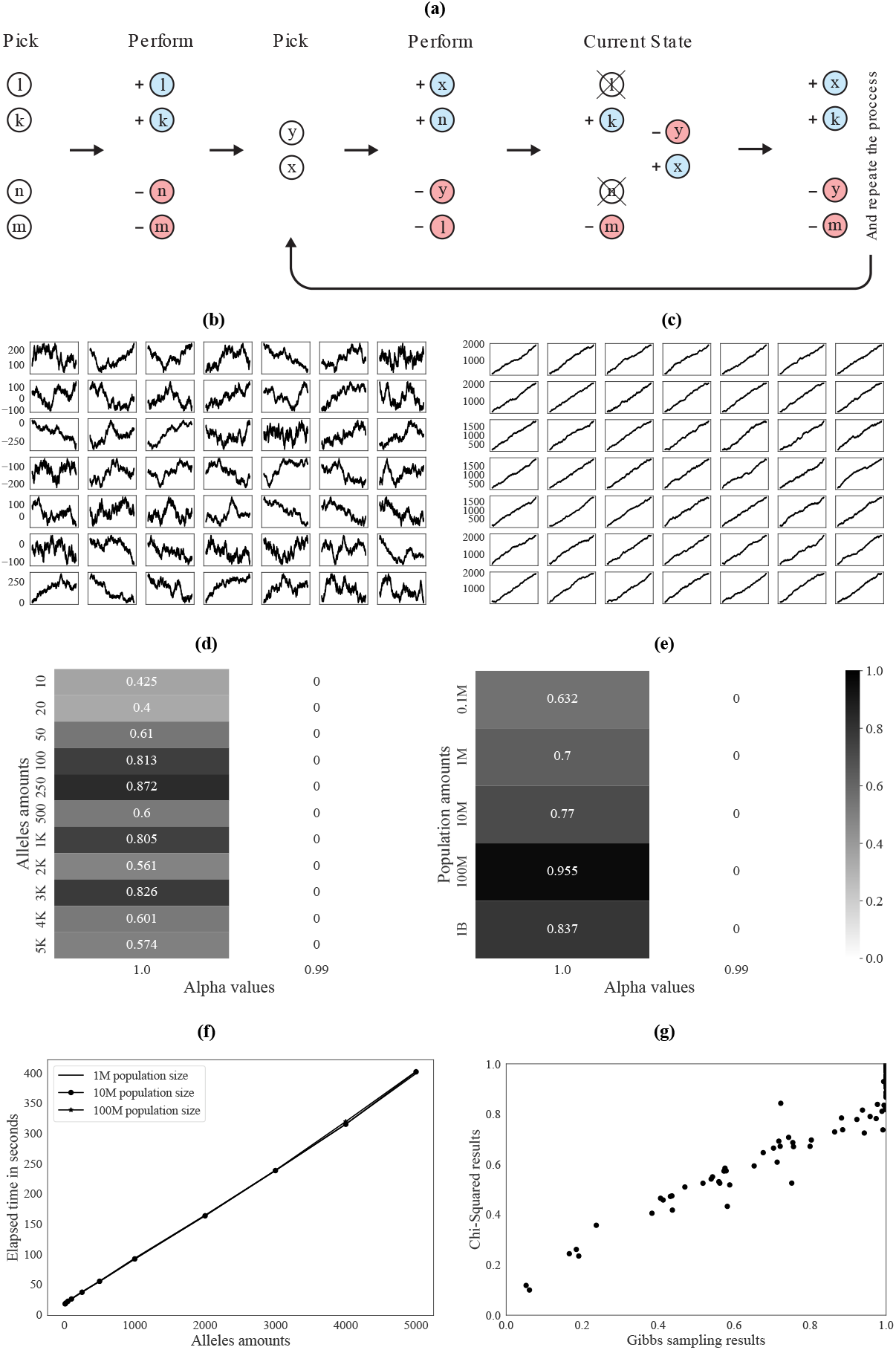
DC case results. **(a)**. Visual representation of the swap of 2 alleles in the UMAT test. First sample an allele *i* according to the alleles distribution, then sample an allele *j* according to the allele pairs distribution given allele *i* and subtract one individual with this pair (it must exist). then, sample a pair of alleles {*k, l*} according to the HWE alleles distribution and add one individual to it. Then, in each iteration, sample an allele *m* according to the allele pairs distribution given allele *l* and subtract one individual from the pair {*m, l*}. Sample an allele *t* according to the allele marginal distribution and add one individual to the pair {*t, j*}. This way, only the marginal observations of the alleles {*t, k, m, i*} are changed. *t* and *k* have one excessive marginal observation, as *l* and *k* had in the beginning and therefore take their place. Note that we can randomly invert their order. Similarly, *m* and *i* are missing one marginal observation, as *j* and *i* had in the beginning and therefore take their place in the next iteration. **(b)**. 49 implementations of UMAT using the same observations in HWE, with *α* = 1.0 (0 ≤ *α* ≤ 1 represents the fit of the simulation to the HWE, as described in Materials and methods), *N* = 1.*e*6 and 1, 000 alleles. **(c)**. 49 implementations of UMAT using the same observations with a slight deviation of HWE, with *α* = 0.99, *N* = 1.*e*6 and 1, 000 alleles. **(d)**. p-value results of UMAT for different allele numbers and alpha values, with *N* = 1.*e*8. **(e)**. p-value results of UMAT for different population sizes and alpha values, with 100 alleles. **(f)**. Elapsed time in seconds for running UMAT using different allele numbers and population sizes, with *α* = 1.0. As one can see, the population size has no effect on the run time. **(g)**. Scatter plot showing for 100 randomly chosen SNPs, the p-value results obtained with Chi-Square and UMAT.

### DC model validation

To test the accuracy of the DC model, we used simulated multi-allele samples and real-world SNP data (See methods).

The simulation was based on the generation of allele pairs using the HWE assumption, and with a probability of 1 − *α* choosing homozygotes instead of the current choice. We tested simulations with different population sizes between *N* = 1, 000 and *N* = 1.*e*8, with 10 − 5, 000 different alleles, and *α* = 0.92, 0.99, 1.0. We first simulated a population of *N* = 1.*e*6 samples with 1, 000 different alleles in HWE (*α* = 1) and with a slight deviation *α* = 0.99. We performed 49 (7 by 7 plot) implementations of UMAT (Fig 1 B vs C). In HWE, the change in the log-likelihood is a random walk around 0 (B plot). In contrast, even a very small deviation of HWE (C plot) shows a consistent increase in the log-likelihood. To show that the results are not sensitive to the population size and allele number, we repeated the analysis with different allele numbers (Fig 1 D) and population size (Fig 1 E). In both cases, the fraction of time steps below the current likelihood is above 0.05 for *α* = 0, and lower than 1.*e* − 3 for *α* = 0.99. We repeated the analysis for both smaller populations and different allele numbers (S1 Fig A, B for *α* = 1.0, 0.92 respectively).

We tested the run time of UMAT. This run time is only based on the need to choose the alleles, which is linear in the number of alleles, and independent of the population size (Fig 1 F).

We further tested the model on bi-allelic SNP data from the 1K genome project. The population may deviate from HWE. We analyzed 100 SNPs and compared the results of UMAT and a classical Chi-Square (given the large population size and the low number of alleles). One can see an excellent fit between the p-value of UMAT and of the Chi-Square (Fig 1 G). However, in general, the Chi-square gives a slight overestimate of *p*.

### Goodness of fit test for AC

We then extended the analysis to ambiguous genetic typing, where the precise allele pair is unknown. For the sake of simplicity, we first described the deviation from the goodness of fit test when there is ambiguity in general, and only then discuss the HWE case. Assume *N* samples and a set of categories, each with an unknown probability *q*_*i*_. We further assumed that given a “real” category *i*, the observation is a distribution over all possible categories *η*_*i*_(*j*)*∀j*, where *η*_*i*_(*j*) is the probability of observing *j* given the real category *i*. The event of having *k* cases of *i* (although we may not observe them given the ambiguity) is a binomial random variable: 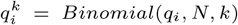 with expectation and variance (for small enough *q*_*i*_) of: *E*_*i*_(*k*) = *N · q*_*i*_, *V*_*i*_(*k*) = *N · q*_*i*_ However, this is not the observed variable. We may observe *j* instead of *i*. Assuming a set of samples, we can define a variable *z*_*j*_ - the total mass of the observation *j. z*_*j*_ is a random variable defined as a sum over donors and alleles:

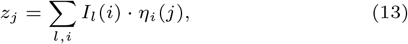

where *I*_*l*_(*i*) is the indicator of event *i* in donor *l. i* is the real allele, and *j* is the observed allele. We can compact the sum over donors to:

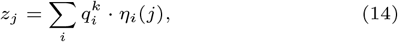

The random variable *z*_*j*_ is a weighted sum of binomials, where we have replaced a multinomial with a set of binomials. This is an approximation, since their sum may be different than *N*, but it is accurate for large enough values of *N*. The expected values of *z*_*j*_ are as before:

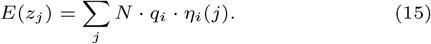

However, in contrast with the classical chi-square, the variances are smaller or equal to the mean:

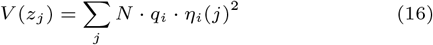

Finally, we assumed that *N* is large enough for the normal distribution approximation on *j* (not on *i*) to hold. In the deterministic case that *η*_*i*_(*j*) = *δ*_*i,j*_, this converges to the classical Chi-Square test. Otherwise, per definition, the variance is smaller.

### Allele pairs

Assume now pairs of alleles *l, k*, and observed allele pairs *m, n. i* above is replaced by *l, k*, and *j* above is replaced by *m, n*. In this case. We can define

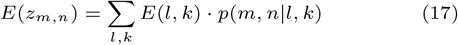

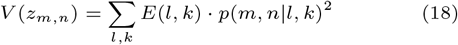

However, we do not know *p*(*m, n*|*l, k*). We thus use a simplifying assumption that *p*(*m, n*|*l, k*) ∼ *p*(*m, n*), and can estimate:

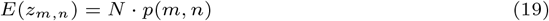

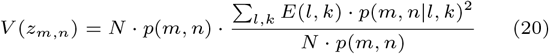

We can thus define a correction to the variance estimate, as a sum over samples *t*:

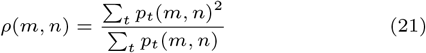

and using the same approximation obtain:

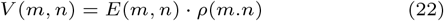

Finally, we revised the goodness of fit test to be:

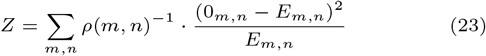

In the absence of ambiguity, *ρ*(*m, n*) is simply 1. Note that the sum contains only non-negative values. if *m, n* is never observed, then there is no correction, and arbitrarily: *ρ*(*m, n*) = 1

### Validation of AC method

Two approaches can be proposed to test the HWE assumption. First, one can sample the distribution of allele pairs in each individual to produce a single pair according to the probability of each candidate pair. Assume for example that a given sample has two candidate pairs with probabilities of 0.3 and 0.7 respectively. One can choose the first with a probability of 0.3 or the second with a probability of 0.7. One can then apply UMAT. To test the sampling, we used the simulation presented above in the DC case and added ambiguity by assigning a probability per sample to replace the real pair with a set of probabilities for possible pairs (See methods). Again, the test is highly accurate with practically no FP and a very accurate detection even for a very small deviation from HWE (Fig 2 A).

**Fig. 2.**
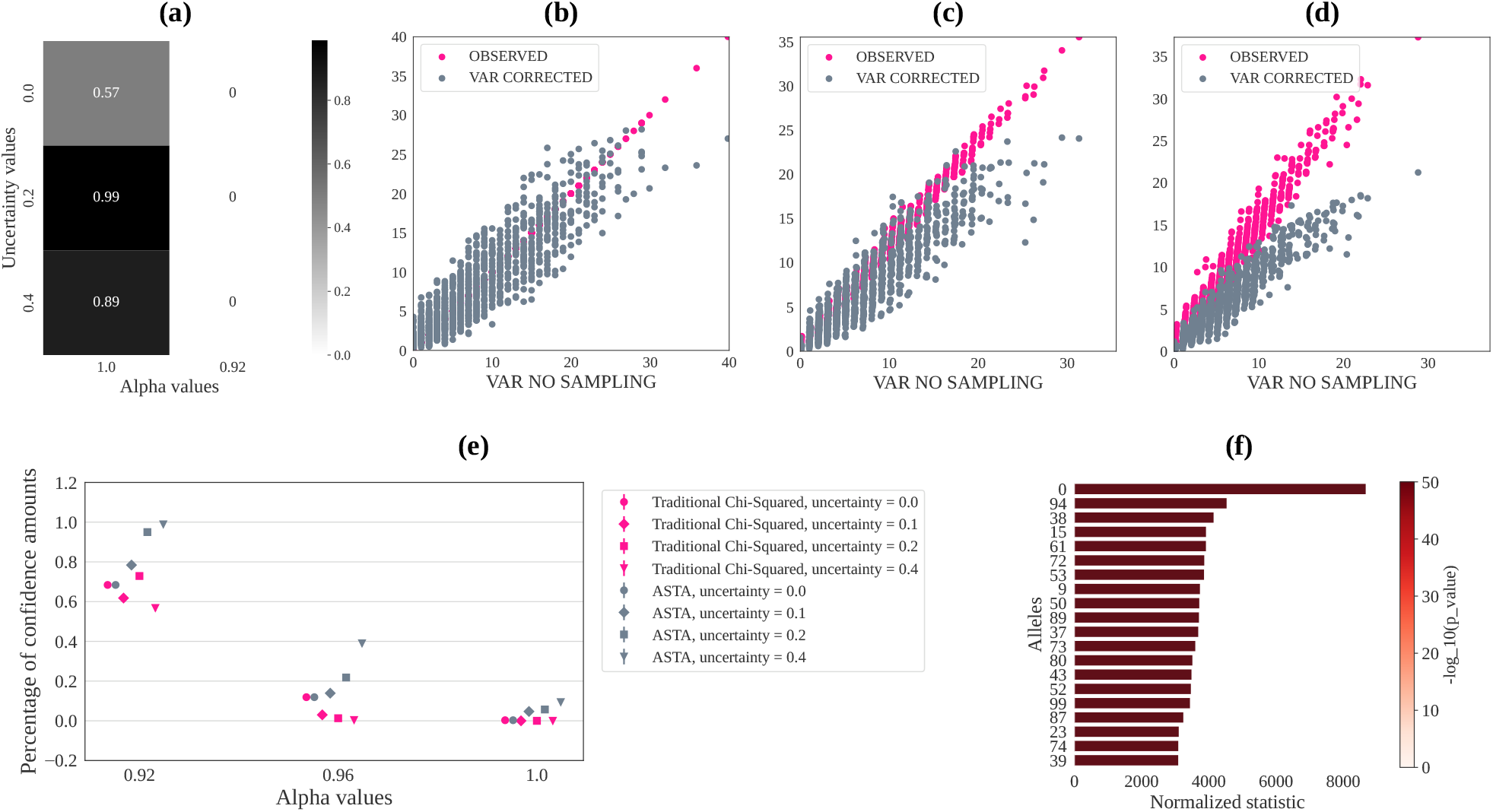
AC case results. **(a)**. p-value results of UMAT with ambiguity for different *ρ* values (*ρ* represents the probability for a sample to have ambiguous typing, as described in Materials and methods) and *α* values, with 100 alleles and *N* = 10, 000. **(b)**-**(d)**. Scatter plot of real vs estimated variance for simulated data with 50 alleles, *N* = 1.*e*4, *α* = 1, *ρ* = 0, 0.2, 0.4 (respectively). Each dot is an allele pair {*i, j*}. The x-axis represents the sampled variance over {*O*_*k*_(*i, j*)}_*k*_, the y-axis represents either the value of *O*(*i, j*) or the corrected denominator used in ASTA: *E*(*i, j*) *· ρ*(*i, j*). **(e)**. Fraction of positive results for *α* = 0.92, 0.96, 1.0, *ρ* = 0.0, 0.1, 0.2, 0.4 and each test: Traditional Chi-Squared, ASTA and Chi-Squared with sampling (i.e. for each person sampling certain alleles given all his possible allele pair observations and then using a traditional Chi-Squared). The results are out of 300 simulations with 5 alleles and *N* = 2000. **(f)**. We simulated data (see Methods) using 100 alleles, *N* = 10, 000, *α* = 1.0. Here all the data is in HWE, except from the pairs containing the 0 allele (top bar). Each bar corresponds to a specific allele and shows the normalized statistic (the sum of Chi-Square term with correction only on pairs containing this allele, and divided by the Degree of Freedom (DOF): the number of pairs containing this allele minus 1), as well as the *−log*(*p value*) for this allele, here the *p value* is calculated using the statistic and DOF of the allele.

An alternative method is to improve the variance estimate in the goodness of fit test. To test the deviation of the variance from the binomial estimate, we compared the variance and the estimated value for each allele pair (*m, n*) as a function of the ambiguity (Fig 2 B-D). As expected, when *α* = 1, the corrected variance estimate is equivalent to the observed value and is approximately the expected value. However, when the ambiguity is increased, the variance is much lower than the expected value but is similar to the corrected variance estimate.

We further tested the regular Chi-square and ASTA for *α* = 0.92 *−* 1.0 and four different values of *ρ* (0.0, 0.1, 0.2, 0.4), we ran the simulation 300 times and averaged the fraction of significance results (each result is either 1 for significance or 0) at the *p* = 0.05 level. We used a population of size *N*_0_ = 2000 and 5 alleles (Fig 2 E).

Both the old Chi-square and ASTA coincide for *ρ* = 0.0 since the term of *V* (*i, j*) is the same in the denominator. Also, as *α* approaches 1.0 (i.e. original observations get closer to HWE), and as *ρ* gets higher, the classical Chi-Square reports a deviation from HWE, while the real observations are in HWE. ASTA does not produce more false positives than randomly expected. Moreover, it detects much more precisely the deviation from HWE, even for small deviations as was the case for UMAT. Results for a higher number of alleles and larger populations are even more accurate and are given in the Supp. Mat. (S2 Fig).

A clear advantage, of this model is the possibility to detect the alleles associated with the deviation from HWE. Instead of summing over all allele pairs, one can sum the Chi-Square term only on pairs containing allele *t*. To test that we performed a simulation, where only pairs containing a given allele are not in HWE. For pairs containing this allele, we choose the HWE distribution with probability *α*, and choose the homozygotes otherwise. To test that we set *α* = 0.7 and perform for every allele *j*:

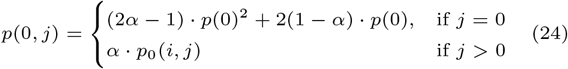

We then tested whether the deviation from HWE was detected and whether the appropriate allele was detected (Fig 2 F). One can see that the main deviation is for the chosen allele (top row). The same holds for simulations with different numbers of alleles and population sizes.

### Deviation of HWE in HLA

To test the deviation from HWE in a large-scale human population, with a large number of alleles, we analyzed the HLA allele frequency distribution from 8.1 million US donors [9]. The HLA locus has 5 classical loci (A,B,C,DRB1,DQB1), and the population is divided into 5 broad populations (CAU - White/European, HIS-Hispanic or Latino, AFA-Black/African American, API - Asian or Pacific Islander and NAM - Native American). Each broad population is further divided into sub-populations. We computed for each broad population and each locus the alleles that deviate the most (highest ASTA score) from HWE (Fig 3). This measurement reproduces multiple previous results. For example, in the CAU populations, the alleles that deviate the most are related to an Ashkenazi Jew haplotyope - *A*26:01 B*38:01 C*12:03 DQB1*03:02 DRB1*04:02* [17]. Another example would be the *A*34:01* that has a clear north-south separation in the Asian population [23]. The top A allele (*A*01:03*) and top B allele (*B*41:01*) in AFA are both from eastern and northern Africa and not sub-Saharan African. *DRB1*04:07* the most out of HWE DRB1 in the AFA population. However, this is a Columbian allele that has been mixed with the AFA population, mainly in Columbia [32; 9].

**Fig. 3.**
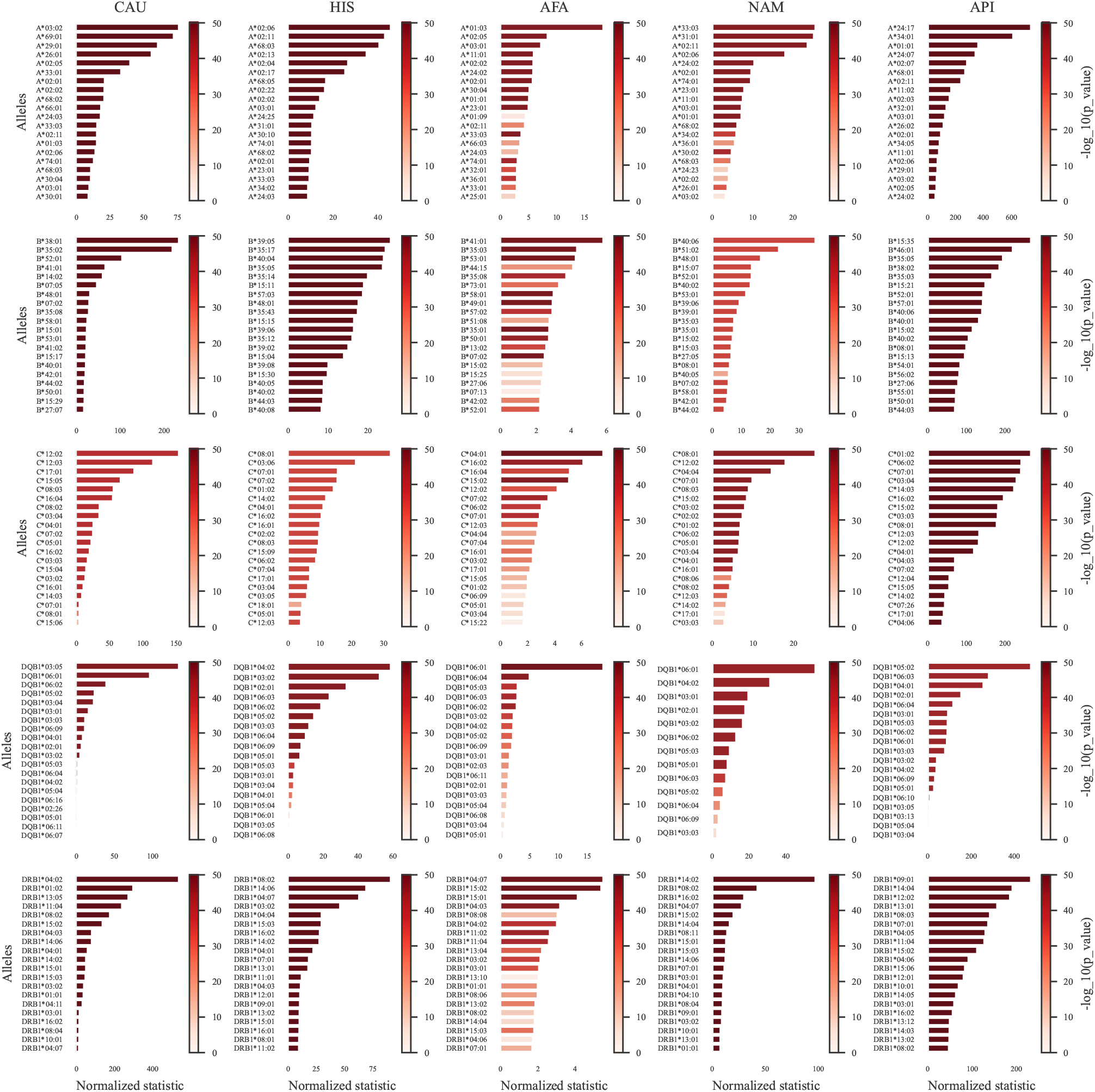
Deviation of alleles from HWE. Each row is one of the 5 Broad US populations. Each column is a locus (A,B,C,DQB1, DRB1). Within each subplot the bar is the ASTA score divided by the DOF. The color represents the log p value. Deeper colors are more significant. The bars are ordered by decreasing significance. Note that the scale of each plot is very different, but the coloring of the bar is consistent among all subplots.

**Fig. 4.**
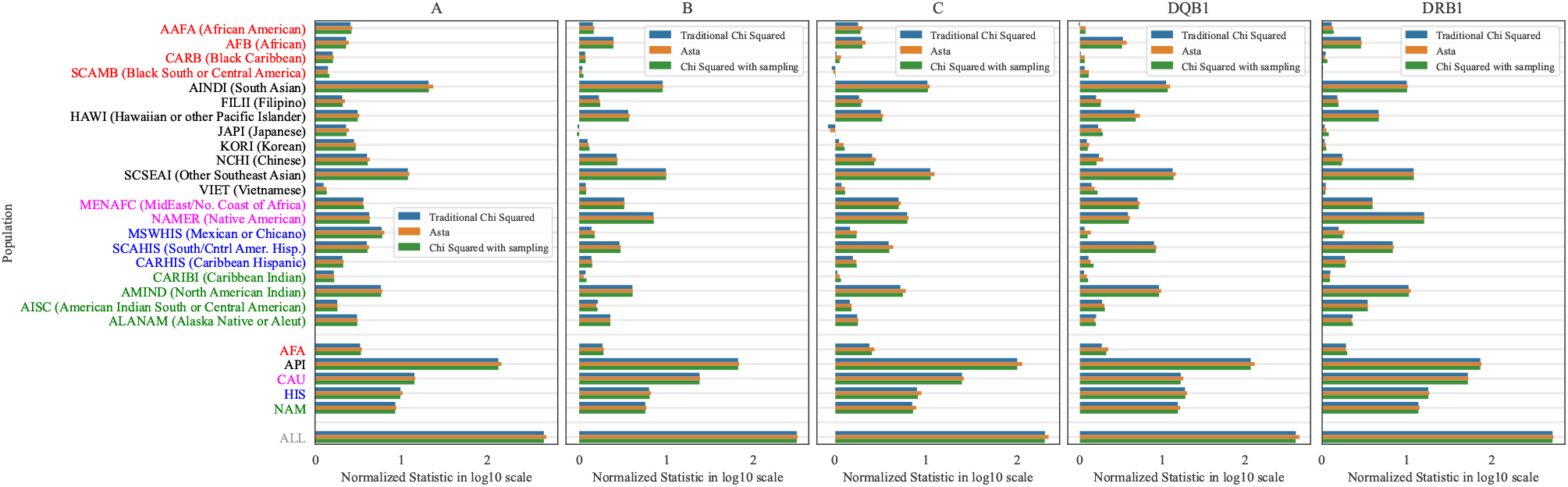
Each subplot is a different locus (A,B,C,DQB1, DRB1). The different colors represent the 10 base log of: the scores divided by the DOF. The ASTA score (Orange) is slightly higher than the classical Chi-Square (Blue) and from the sampling (Green). Note that in this case, the ambiguity is limited. As such, the differences are not very large. The top populations are the detailed population, followed by the broad populations, followed by the entire donor registry. Each detailed population is colored according to the broad population.

**Fig. 5.**
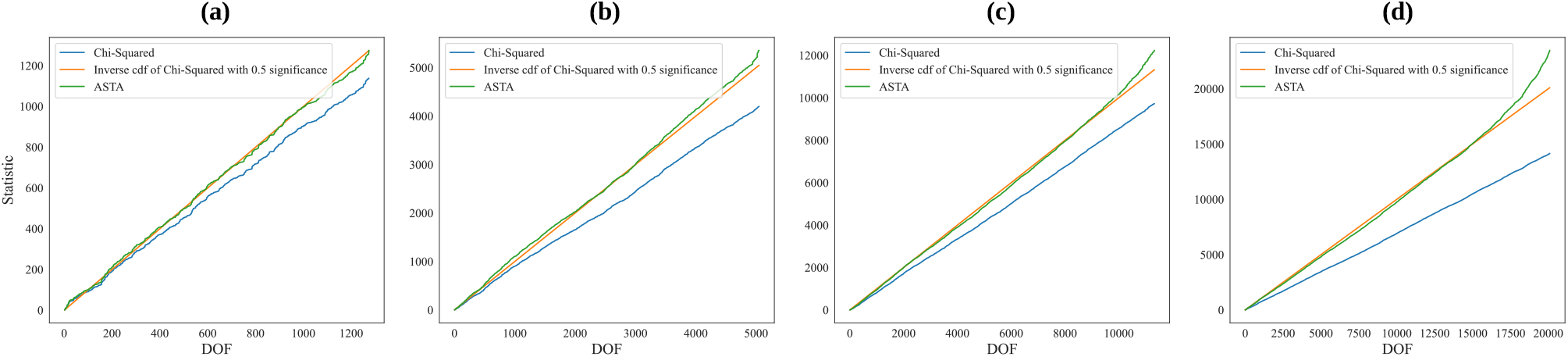
A comparison between: Chi-Squared, ASTA and Chi-Squared with sampling. **(a)**. 50 alleles, 50, 000 population size, *α* = 0.1, *ρ* = 0.1. **(b)**. 100 alleles, 200, 000 population size, *α* = 0.1, *ρ* = 0.2. **(c)**. 150 alleles, 250, 000 population size, *α* = 0.1, *ρ* = 0.15. **(d)**. 200 alleles, 400, 000 population size, *α* = 0.1, *ρ* = 0.3.

To further study the total deviation from HWE in the total population, we compared all three models (AST, Chi-Square, Chi-Square with sampling) on all broad and detailed groups and for all loci. The population with the strongest deviation from HWE is the Asian Pacific population (API), since it includes a combination of multiple sub-populations, including Chinese and Indian. Within it the API population the Indian (AINDI) and mixed Asian (SCSEAI) have the highest deviation in all loci. The homogeneous populations, like Japan, Korea and Vietnam have consistently very low deviation from HWE. Interestingly, the Mexican population (MSWHIS) deviates the most in the A locus.

## Supporting information

**S1 Table**.

**Population codes**. NMDP US population codes, composed of 21 detailed populations, group into 5 broad populations..

**S1 Fig**.

**UMAT results for small populations**. p-value results for different allele numbers and population amounts, with: **(a)**. *α* = 1.0. **(b)**. *α* = 0.92.

**S2 Fig**.

**Comparison between ASTA and Traditional Chi-Squared**. Fraction of positive results for *α* = 0.92, 0.96, 1.0, *ρ* = 0.0, 0.1, 0.2, 0.4 and each test: Traditional Chi-Squared, ASTA and Chi-Squared with sampling (i.e. for each person sampling certain alleles given all his possible allele pair observations and then using a traditional Chi-Squared). The results are out of 300 simulations with 5 alleles and *N* = 2000. For each subplot we used different population amounts and allele numbers: **(a)**. 50 alleles, *N* = 50, 000. **(b)**. 100 alleles, *N* = 200, 000. **(c)**. 150 alleles, *N* = 250, 000. **(d)**. 200 alleles, *N* = 400, 000.

**S3 Fig**.

**Deviation of alleles from HWE**. Each row is one of the 21 US populations. Each column is a locus (A,B,C,DQB1, DRB1). Within each subplot the bar is the ASTA score divided by the DOF. The color represents the log p value. Deeper colors are more significant. The bars are ordered by decreasing significance. Note that the scale of each plot is very different, but the coloring of the bar is consistent among all subplots.

## Conclusion

The HWE assumption is crucial for many population genetics models including, among many others Expectation Maximization (EM) based haplotype frequency estimate [18; 16], Imputation, admixture algorithms [15; 4; 26], and any model containing an estimate of probability of allele pairs. We have here focused on the specific context of multi-allelic loci in diploids. We propose in this context two algorithms for the estimate of deviation from HWE. The first (UMAT) is a method to test the deviation of the exact distribution from the one expected in HWE, and the second (ASTA) is a goodness of fit test that can handle ambiguous typing. UMAT is based on a perturbative approach, where we test the effect of randomly swapping pairs of alleles on the likelihood. ASTA is based on the correction of the Chi-Square test to reflect the variance with ambiguous typing. We show the accuracy of both models using extensive simulation and real-world cases. ASTA is per definition an approximation, but it has the advantage of detecting the alleles/haplotypes most associated with the deviation from HWE.

We have tested both methods using the well-known HLA locus in the human chromosome 6, and shown a significant deviation from HWE in practically all genes and all sub-populations, with a larger deviation, in broad populations than in detailed populations. We further detected the alleles most associated with deviation from HWE, and have shown it reproduces multiple previous reports on deviations from HWE in HLA.

The main advantage of UMAT and ASTA over existing methods is the possibility of estimating the deviation from HWE from thousands or tens of thousands of alleles/haplotypes, and with no limit on the population size. Both methods have limitations that are handled by other models. Those include among others the limitation to a single locus and the lack of a solution for a structured population [13]. In the case of ASTA, there are other limitations, including the normal approximation, which is only valid if we sum enough elements. Note that for ambiguous typing, the total number of elements in the sum can be much larger than the expected value. To handle that, we allow the user to limit the test to a minimal denominator, and ignore allele/haplotype pairs with too low denominators. Another quite stringent assumption is that the expectation of the observed value is indeed the allele pair probability. This is trivial in the regular goodness of fit test. However, in the ambiguous case, the ambiguity may be biased toward specific pairs. We currently have no solution if this assumption is not at least approximately valid.

## Key Points

- Hardy Weinberg Equilibrium estimate is crucial for many bioinformatics tools. However, there is no accurate estimator for the HWE in large multi-locus populations.
- We present two algorithms to solve that. UMAT based on GIBBS sampling, and a corrected goodness of fit algorithm named ASTA.
- UMAT can detect very small deviations from HWE in a short time even with a very large number of alleles and in large populations.
- ASTA can further detect the alleles associated with deviation from HWE.
- We applied those to an 8 million sample of HLA genotypes, and detected novel alleles associated with deviation from HWE.

## Funding

The work of MM and YL was partially funded by the Office of Naval Research grant (N00014-23-1-2057) (https://govtribe.com/vendors/national-marrow-donor-program1msy6). The work of SI was funded by ISF grant 870/20 (https://www.isf.org.il/#/) and by a DSI Grant (https://dsi.biu.ac.il/).

## Data availability statement

The code used for simulations is available at https://github.com/louzounlab/HWE_Simulations. The SNP data is available at KAGGLE: https://www.kaggle.com/datasets/extraflash/snipping-data. The HLA frequencies data is availabale at

https://www.allelefrequencies.net/. The HLA genotypes are available upon request from Martin Maiers at the NMDP.

## Competing interests

No competing interest is declared.

## Acknowledgments

The authors thank the anonymous reviewers for their valuable suggestions. We thank Miriam Beller for the English editing.

## Sapir Israeli

Sapir Israeli is a PhD candidate in the Department of Mathematics at Bar Ilan University in Israel, earned dual bachelor’s degrees in Computer Science and Neuroscience, and a master’s degree in Computer Science. Her research integrates machine learning techniques with the study of Human Leukocyte Antigen (HLA) and its applications in Stem cell transplants.

## Yoram Louzoun

Yoram Louzoun is a Professor of Mathematics at the Bar Ilan University in Israel. He has graduated from the Hebrew University in 2000 and continued to a post-doc in Princeton. In 2002, he started a position at Bar Ilan in the Mathematics department. He studies stochastic processes and machine learning, with a special focus on its application to immunology, transplants and microbiome. He has developed a set of algorithms to optimize Stem cells transplants, and machine learning algorithms to predict the host condition from his microbiome or from his T cell repertoire, as well as graph based algorithms.

## Martin Maiers

Martin Maiers is Vice President of Research at Be The Match where he has worked for 28 years. He leads an R&D translation program focused on applying new tools for HLA matching and algorithm development to improve cell and gene therapies. With his team he has developed a number of tools and methods for searching large donor registries with missing or partial information to identify suitable hematopoietic stem cell donors. He holds a degree in Mathematics from the University of Wisconsin and a Master’s Degree in Computational Biology from the University of Minnesota.

## Or Shkuri

Or Shkuri is a MSc student in the Department of Mathematics at Bar Ilan University in Israel, who earned a bachelor’s degree in Applied Mathematics. He currently researches in the area of Machine Learning and Statistics, with an emphasis on Machine Learning with missing features.

## Yuli Tshuva

Yuli Tshuva is a BSc student in the Department of Mathematics at Bar Ilan University in Israel. He currently researches in the area of Machine Learning in the YOLO lab at Bar Ilan University for prof. Yoram Louzoun.

## Notes

https://www.kaggle.com/datasets/extraflash/snipping-data

https://www.allelefrequencies.net/

### Competing Interest Statement

The authors have declared no competing interest.

### Summary of Updates

Fixed some typos and overall changed the template of the paper to the OUP template

