## Supplementary material for "Efficient test for deviation from Hardy Weinberg Equilibrium with known or ambiguous typing in highly polymorphic loci": S1 Table.

| Detailed population Code | Broad population | Detailed population description |
|---|---|---|
| AAFA | AFA - Afro Americans | African American |
| AFB | | African |
| CARB | | Black Caribbean |
| SCAMB | | Black South or Central America |
| AINDI | API - Asian or Pacific Islands | South Asian |
| FILII | | Filipino |
| HAWI | | Hawaiian or other Pacific Islander |
| JAPI | | Japanese |
| KORI | | Korean |
| NCHI | | Chinese |
| SCSEAI | | Other Southeast Asian |
| VIET | | Vietnamese |
| NAMER | CAU - Caucasian | North American White |
| MENAFC | | MidEast/No. Coast of Africa |
| MSWHIS | HIS - Hispanic | Mexican or Chicano |
| SCAHIS | | South/Cntrl Amer. Hisp. |
| CARHIS | | Caribbean Hispanic |
| CARIBI | NAM - Native Americans | Caribbean Indian |
| AMIND | | North American Indian |
| AISC | | American Indian South or Central American |
| ALANAM | | Alaska Native or Aleut |

Table S1: NMDP US population codes, composed of 21 detailed populations, group into 5 broad populations.
