## Supplementary figures and images for "Efficient test for deviation from Hardy Weinberg Equilibrium with known or ambiguous typing in highly polymorphic loci"

### S1 Fig.

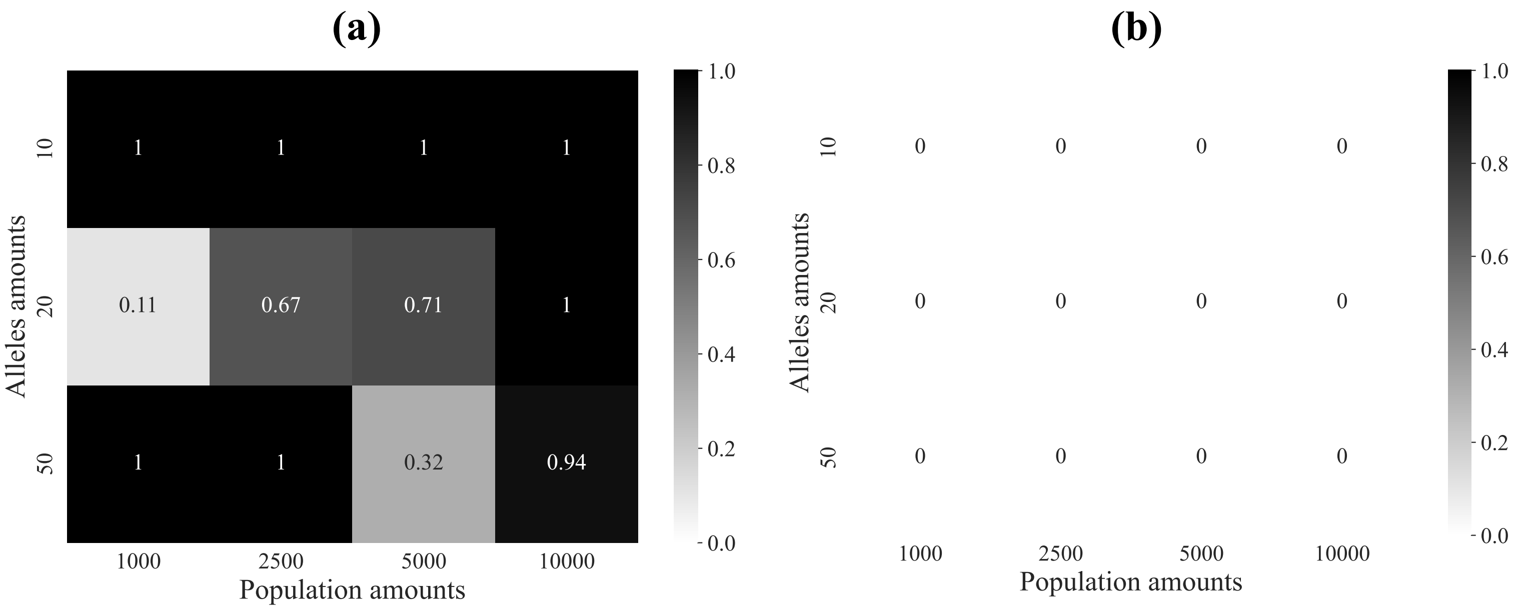

### S2 Fig.

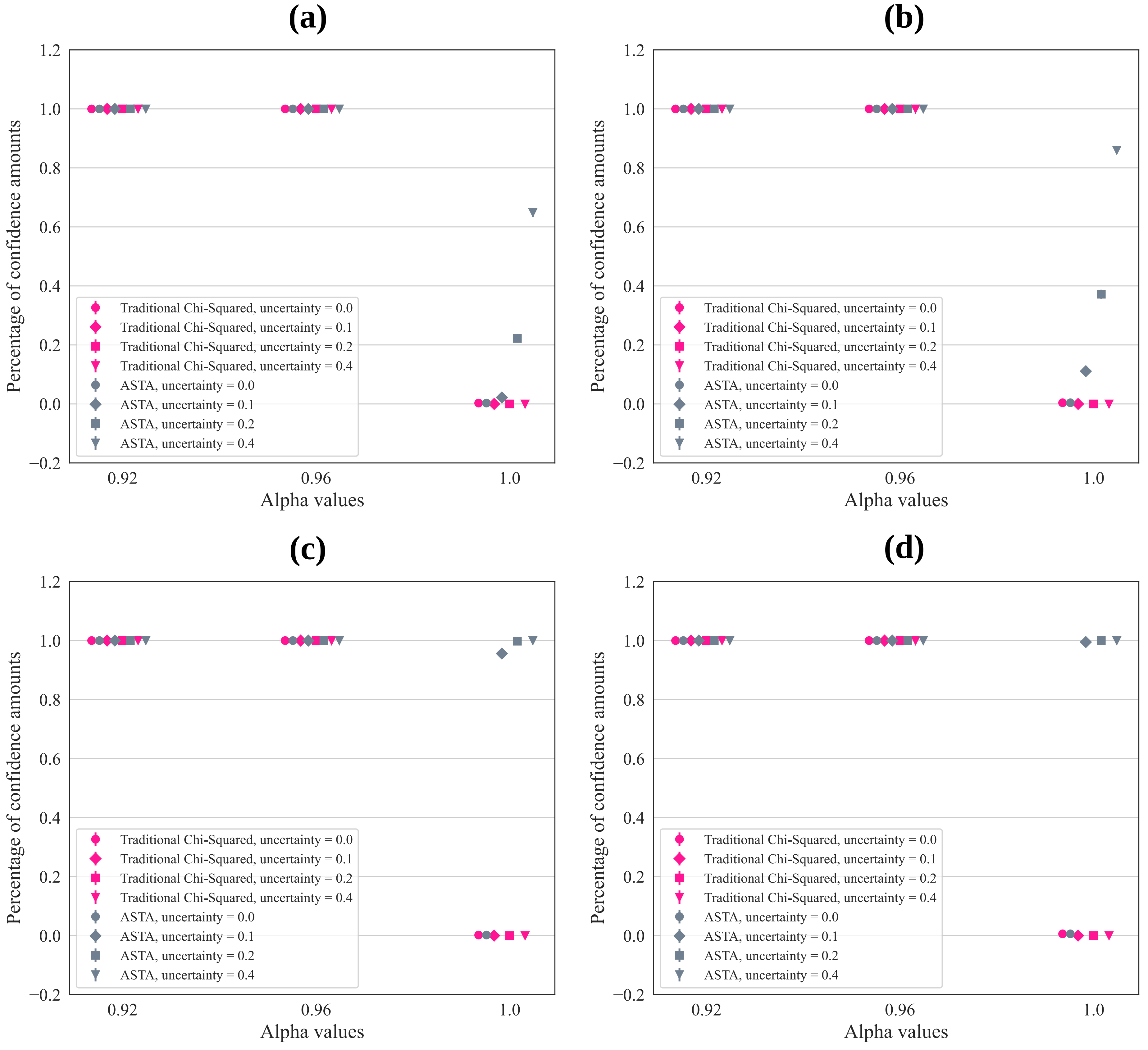

### S3 Fig.

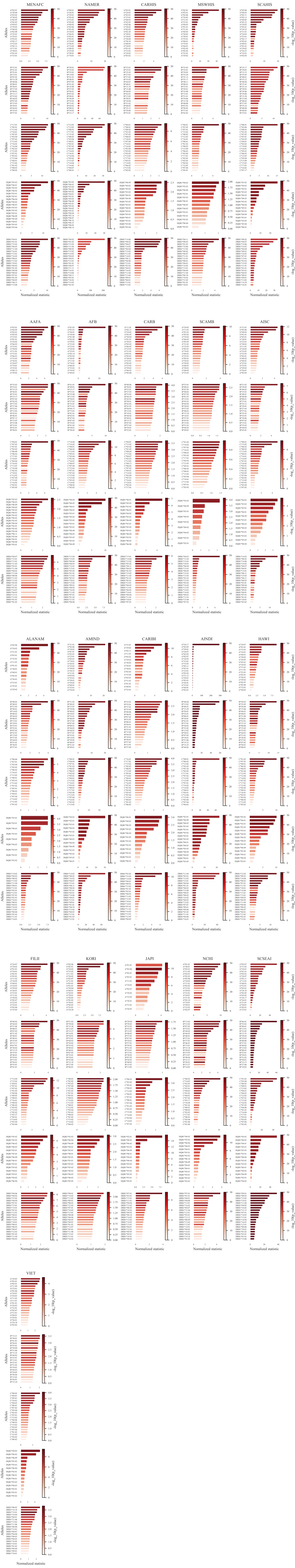
